# Motility and Phototaxis of *Gonium*, the Simplest Differentiated Colonial Alga

**DOI:** 10.1101/845891

**Authors:** Hélène de Maleprade, Frédéric Moisy, Takuji Ishikawa, Raymond E. Goldstein

**Affiliations:** Department of Applied Mathematics and Theoretical Physics, Centre for Mathematical Sciences, University of Cambridge, Wilberforce Road, Cambridge CB3 0WA, United Kingdom; Laboratoire FAST, Université Paris-Sud, CNRS, Universtié Paris-Saclay - Paris, France; Department of Finemechanics, Graduate School of Engineering, Tohoku University, 6-6-01 Aoba, Aramaki, Aoba-ku, Sendai 980-8579, Japan

## Abstract

Green algae of the *Volvocine* lineage, spanning from unicellular *Chlamydomonas* to vastly larger *Volvox*, are models for the study of the evolution of multicellularity, flagellar dynamics, and developmental processes. Phototactic steering in these organisms occurs without a central nervous system, driven solely by the response of individual cells. All such algae spin about a body-fixed axis as they swim; directional photosensors on each cell thus receive periodic signals when that axis is not aligned with the light. The flagella of *Chlamydomonas* and *Volvox* both exhibit an adaptive response to such signals in a manner that allows for accurate phototaxis, but in the former the two flagella have distinct responses, while the thousands of flagella on the surface of spherical *Volvox* colonies have essentially identical behaviour. The planar 16-cell species *Gonium pectorale* thus presents a conundrum, for its central 4 cells have a *Chlamydomonas*-like beat that provide propulsion normal to the plane, while its 12 peripheral cells generate rotation around the normal through a *Volvox*-like beat. Here, we combine experiment, theory, and computations to reveal how *Gonium*, perhaps the simplest differentiated colonial organism, achieves phototaxis. High-resolution cell tracking, particle image velocimetry of flagellar driven flows, and high-speed imaging of flagella on micropipette-held colonies show how, in the context of a recently introduced model for *Chlamydomonas* phototaxis, an adaptive response of the peripheral cells alone leads to photo-reorientation of the entire colony. The analysis also highlights the importance of local variations in flagellar beat dynamics within a given colony, which can lead to enhanced reorientation dynamics.

## I. INTRODUCTION

Since the work of A. Weismann on germ-plasm theory in biology [1] and of J.S. Huxley on the nature of the individual in evolutionary theory [2], the various species of green algae belonging to the family *Volvocaceae* have been recognized as important ones in the study of evolutionary transitions from uni-to multicellular life. In a modern biological view [3], this significance arises from a number of specific features of these algae, including the fact that they are an extant family (obviating the need to study fossils), are readily obtainable in nature, have been studied from a variety of perspectives (biochemical, developmental, genetic), and have had significant ecological studies. From a fluid dynamical perspective [4], their relatively large size and easy culturing conditions allow for precise studies of their motility, the flows they create with their flagella, and interactions between organisms, while their high degree of symmetry simplifies theoretical descriptions of those same phenomena [5].

As they are photosynthetic, the ability of these algae to execute phototaxis is central to their life. Because the lineage spans from unicellular to large colonial forms, it can be used to study the evolution of multicellular coordination of motility. Motility and phototaxis of motile green algae have been the subjects of an extensive literature in recent years [6–13], focusing primarily on the two extreme cases: unicellular *Chlamydomonas* and much larger *Volvox*, with species composed of 1, 000 – 50,000 cells. *Chlamydomonas*, the simplest member of the *Volvocine* family, swims typically by actuation of its two flagella in a breast stroke, combining propulsion and slow body rotation. It possesses an *eye spot*, a small area highly sensitive to light [14, 15], which triggers the two flagella differently [16]. Those responses are adaptive, on a timescale matched to the rotational period of the cell body [17–19], and allow cells to scan the environment and swim towards light [13]. Multicellular *Volvox* shows a higher level of complexity, with differentiation between interior germ cells and somatic cells dedicated to propulsion. Despite lacking a central nervous system to coordinate its cells, *Volvox* exhibits accurate phototaxis. This is also achieved by an adaptive response to changing light levels, with a response time tuned to the *colony* rotation period which creates a differential response between the light and dark sides of the spheroid [7, 20].

In light of the above, a natural questions is: how does the simplest *differentiated* organism achieve phototaxis? In the Volvocine lineage the species of interest is *Gonium*. This 8- or 16-cell colony represents one of the first steps to true multicellularity [22], presumed to have evolved from the unicellular common ancestor earlier than other Volvocine algae [23]. It is also the first to show cell differentiation. We focus here on 16-cell colonies, which show a higher degree of symmetry than those with 8, but our results apply to both.

A 16-cell *Gonium* colony is shown in Fig. 1. It is organized into two concentric squares of respectively 4 and 12 cells, each biflagellated, held together by an extracellular matrix [24]. All flagella point out on the same side: it exhibits a much lower symmetry than *Volvox*,lacking anterior-posterior symmetry. Yet it performs similar functions to its unicellular and large colonies counterparts as it mixes propulsion and body rotation, and swims efficiently towards light [6, 25, 26]. The flagellar organization of inner and peripheral cells deeply differs [27, 28]: central cells are similar to *Chlamydomonas*, with the two flagella beating in an opposing breast stroke, and contribute mostly to the forward propulsion of the colony. Cells at the periphery, however, have flagella beating in parallel, in a fashion close to *Volvox* cells [21]. This minimizes steric interactions and avoids flagella crossing each other [6]. Moreover, these flagella are implanted with a slight angle and organized in a pin-wheel fashion (see Fig. 1b) [27]: their beating induces a left-handed rotation of the colony, highlighted in Figs. 1c&d and in Supplementary Movie 1. Therefore, the flagella structure of *Gonium* reinforces its key position as intermediate in the evolution towards multicellularity and cell differentiation.

**FIG. 1.**
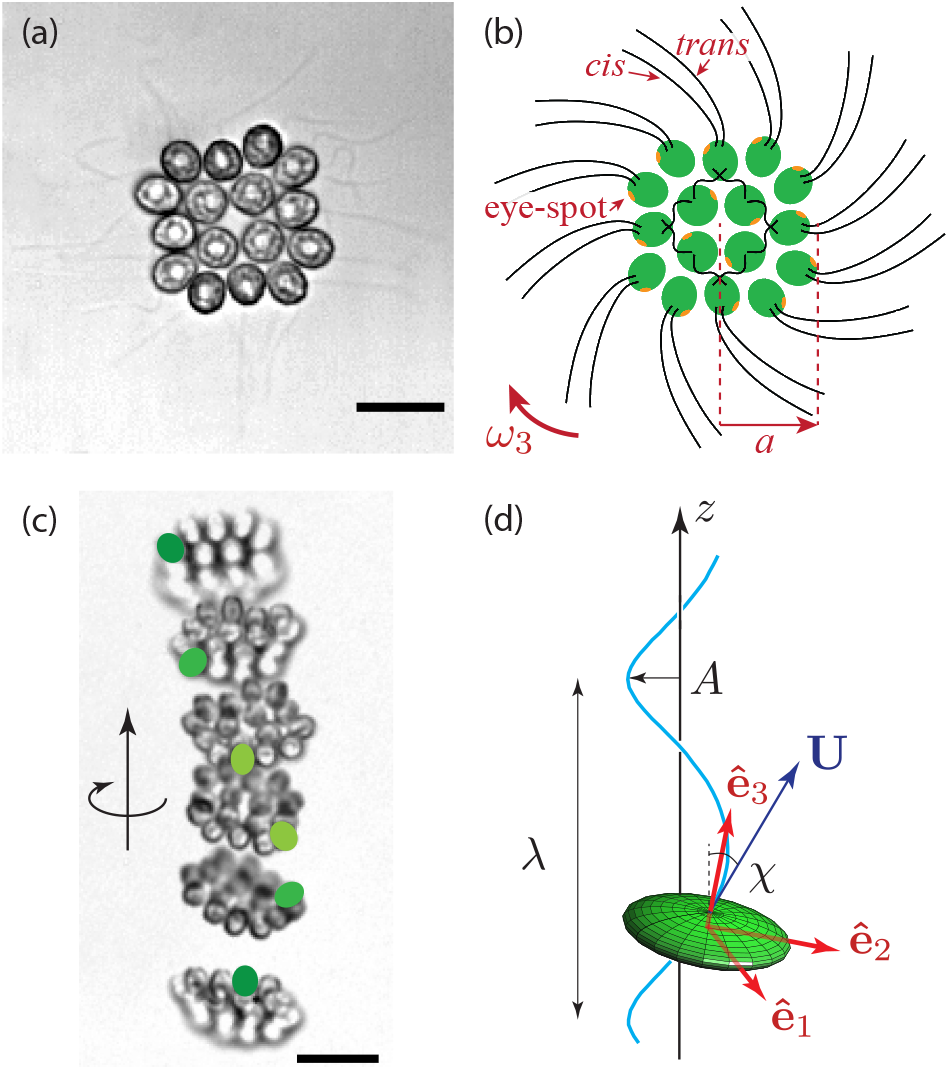
Geometry and locomotion of *Gonium pectorale*. (a) 16-cell colony. Each cell has two flagella, 30 – 40 *μ*m long. Scale bar is 10 *μ*m. (b) Schematic of a colony of radius *a*: 16 cells (green) each with one eye spot (orange dot). The *cis* flagellum is closest to the eye-spot, the *trans* flagellum is furthest [21]. Flagella of the central cells beat in an opposing breaststoke, while the peripheral flagella beat in parallel. The pinwheel organization of the peripheral flagella leads to a left-handed body rotation at a rate *ω*_3_. (c) Upward swimming of a colony. Superimposition of images separated by 0.4 s. Green spots label one specific cell to highlight the lefthanded rotation. Scale bar is 20 *μ*m. (d) Sketch of a helical trajectory: a colony (green ellipsoid) swims with velocity **U** along an oscillatory path (blue line) of wavelength λ, amplitude *A* and pitch angle *χ*. The frame (**ê**_1_, **ê**_2_, **ê**_3_) is attached to the *Gonium* body.

These small flat assemblies show intriguing swimming along helical trajectories - with their body plane almost normal to the swimming direction - that have attracted the attention of naturalists since the eighteenth century [25, 26, 29]. Yet, the way in which *Gonium* colonies bias their swimming towards the light remains unclear. Early microscopic observations have identified differential flagellar activity between the illuminated and the shaded sides of the colony as the source of phototactic reorientation [25, 26]. Yet, a full fluid-dynamics description, quantitatively linking the flagellar response to light variations and the hydrodynamic forces and torques acting on the colony, is still lacking. From an evolutionary perspective, phototaxis in *Gonium* also raises a number of fundamental issues: to what extent is the phototactic strategy of the unicellular ancestor retained in the colonial form? How is the phototactic flagella reaction adapted to the geometry and symmetry of the colony, and how does it lead to an effective reorientation?

Taking this specific structure into account, here we aim to understanding how the individual cell reaction to light leads to reorientation of the whole colony. Any description of phototaxis must build on an understanding of unbiased swimming, so we first focus on the helical swimming of *Gonium*, and show how it results from an uneven distribution of forces around the colony. We then investigate experimentally its phototaxis, by describing the reorientation trajectories and characterising the cells’ response to light. That response is shown to be adaptive, and we therefore extend a previously introduced model for such a response to the geometry of *Gonium*, and show how the characteristic relaxation times are finely tuned to the *Gonium* body shape and rotation rate to perform efficient phototaxis.

## II. FREE SWIMMING OF *GONIUM*

### A. Experimental observations

We recorded trajectories of *Gonium* colonies freely swimming in a sealed chamber on an inverted microscope, connected to a high speed video camera, as sketched in Fig. 2a and detailed in Appendix A. To obtain unbiased random swimming trajectories, we used red light illumination. Trajectories were reconstructed using a standard tracking algorithm, and shown in Fig. 3a and Supplementary Movie 2. They exhibit a large variation in waviness, with some colonies swimming along nearly straight lines, while others show highly curved helices. From Fig. 1c-d, we infer that a colony performs a full body rotation per helix wavelength. This observation suggests that the waviness of the trajectories arises from an uneven distribution of forces developed by the peripheral flagella, with the most active flagella located on the outer side of the helix.

**FIG. 2.**
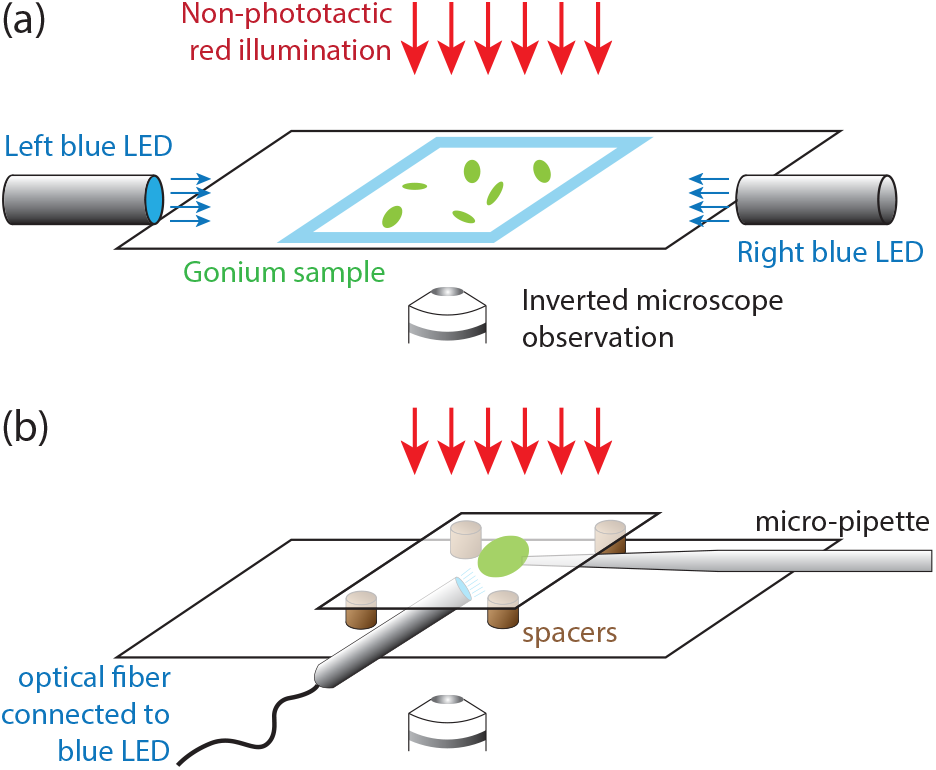
Phototaxis experiments. (a) *Gonium* colonies swim in a sealed chamber made of two glass slides, with nonphototactic red illumination from above, on the stage of an inverted microscope connected to a high speed video camera. Two blue LEDs on the right- and left-hand sides of the chamber can independently shine light with controllable intensities. (b) Micropipette experiments. A micropipette of inner diameter ~ 20 *μ*m holds a colony (green disc) in a chamber made of two glass slides, spaced to allow room for optical fiber connected to a blue LED to enter the chamber.

From image analysis, we extracted for each colony the body radius *a*, the body rotation frequency *ν*_3_ = *ω*_3_/2*π*, the instantaneous swimming velocity *v* projected in the plane of observation, and the mean swimming velocity *v_m_*. The rotation frequency *ν*_3_ ~ 0.4 Hz, decreases slightly with colony size as shown in Fig. 3b. This places *Gonium* in a consistent intermediate position between *Chlamydomonas* and *Volvox*, whose radii are respectively about 5 *μ*m and 200 *μ*m, and whose rotation rates are 2 Hz and 0.2 Hz [7, 30]. With a typical flagellar beating frequency of 10 Hz (see measurements in Sec. III), there are ~ 25 flagella strokes per body rotation, a value similar to that in *Chlamydomonas*.

**FIG. 3.**
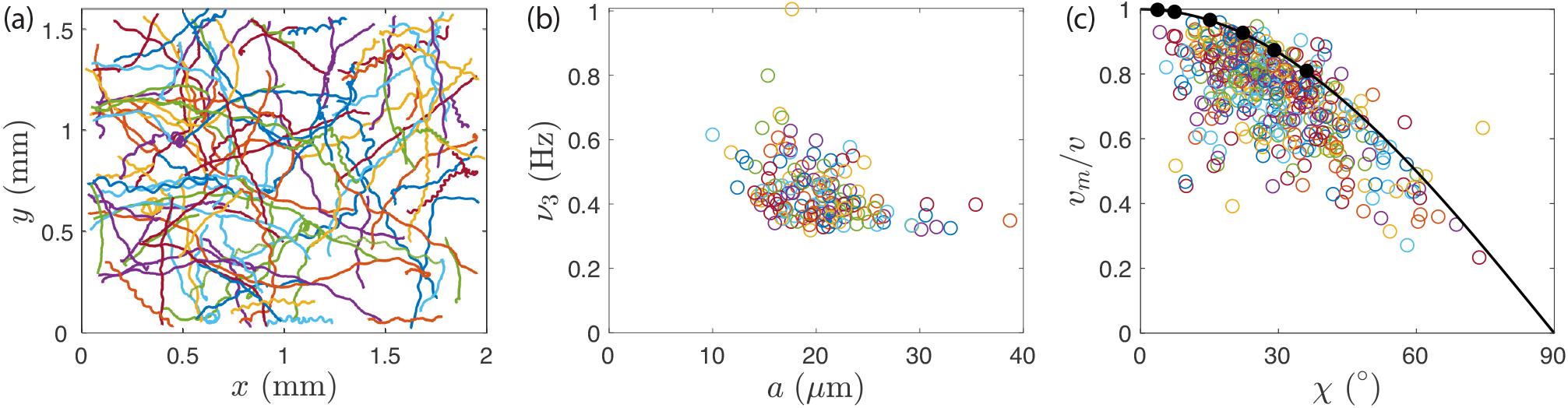
Free swimming of *Gonium*. (a) Trajectories of many colonies under non-phototactic illumination, showing random swimming. Each colored line shows the path of one colony (sample size: 117 colonies). (b) Body rotation frequency *ν*_3_ as a function of colony radius *a* (sample size: 159 colonies). (c) Mean velocity *v_m_* normalised by the instantaneous swimming velocity *v* as a function of helix pitch angle *χ* (sample size: 430 colonies). The black line indicates relation *v_m_*/*v* = cos *χ* and black discs along it are results from the numerical simulations as described in text.

There is a significant correlation between the swimming velocity and the waviness of the trajectories, displayed in Fig. 3c. We quantify the degree of waviness with the pitch angle *χ*, defined as the angle between the swimming speed **U** and the helix axis **ê**_*z*_, such that tan *χ* = (2*π A*/λ), with *A* and λ the helix amplitude and wavelength (see Fig. 1d). Swimming in helices is clearly at the cost of the swimming efficiency: the ratio *v_m_*/*v* shows a marked decrease with the angle *χ*. The data is well described by the simple law *v_m_*/*v* = cos *χ* expected for a helix traced out at constant instantaneous velocity (black line in Fig. 3c). This geometrical law is valid for a 3-dimensional velocity *v*_3*D*_, while we only have access to its two-dimensional projection. In Appendix B we show that *v* (the projected velocity) provides a reasonable approximation to the real velocity *v*_3 *D*_ for *χ* not too large. The average pitch angle in Fig. 3c is *χ* ≃ 30° ± 13°, corresponding to a mean velocity experimentally 30% slower than the instantaneous velocity. In spite of this decreased swimming efficiency, the surprisingly high level of waviness found in most *Gonium* colonies suggest that this trait provides an evolutionary advantage.

### B. Fluid dynamics of the swimming of *Gonium*

We introduce here the fluid dynamical and computational description of the swimming of *Gonium*. With a typical size of 40 *μ*m and swimming speed of 40 *μ*m/s, the Reynolds number is of order 10^−3^, so the swimming is governed by the Stokes equation, *i.e*. by the balance between the force and torque induced by the flagella motion to those arising from viscous drag [31].

We model a colony as a thick disk of symmetry axis **ê**_3_. This disk includes the averaged cell body radius *a*, and also the flagella, which contribute significantly to the total friction. We therefore consider an effective radius *R* encompassing the whole structure. In the frame (**e**_1_, **ê**_2_, **ê**_3_) attached to the body, the viscous force **F**_*v*_ and torque **L**_*v*_ are linearly related to the velocity **U** and angular velocity **Ω** through

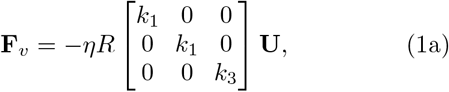

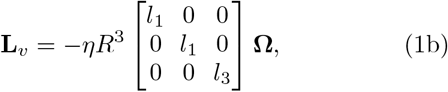

where *η* = 1 mPa s is the viscosity of water, and the numbers (*k*_1_, *k*_3_, *l*_1_, *l*_3_) quantifying the translation and rotation friction along the transverse and axial directions respectively, characteristic of the *Gonium* geometry. Lefthanded body rotation implies **U · Ω** < 0. The angular velocity **Ω** is expressed in the body frame by introducing the Euler angles (*θ, ϕ, ψ*) defined in Fig. 4a [32],

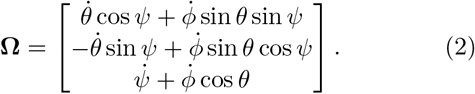

**FIG. 4.**
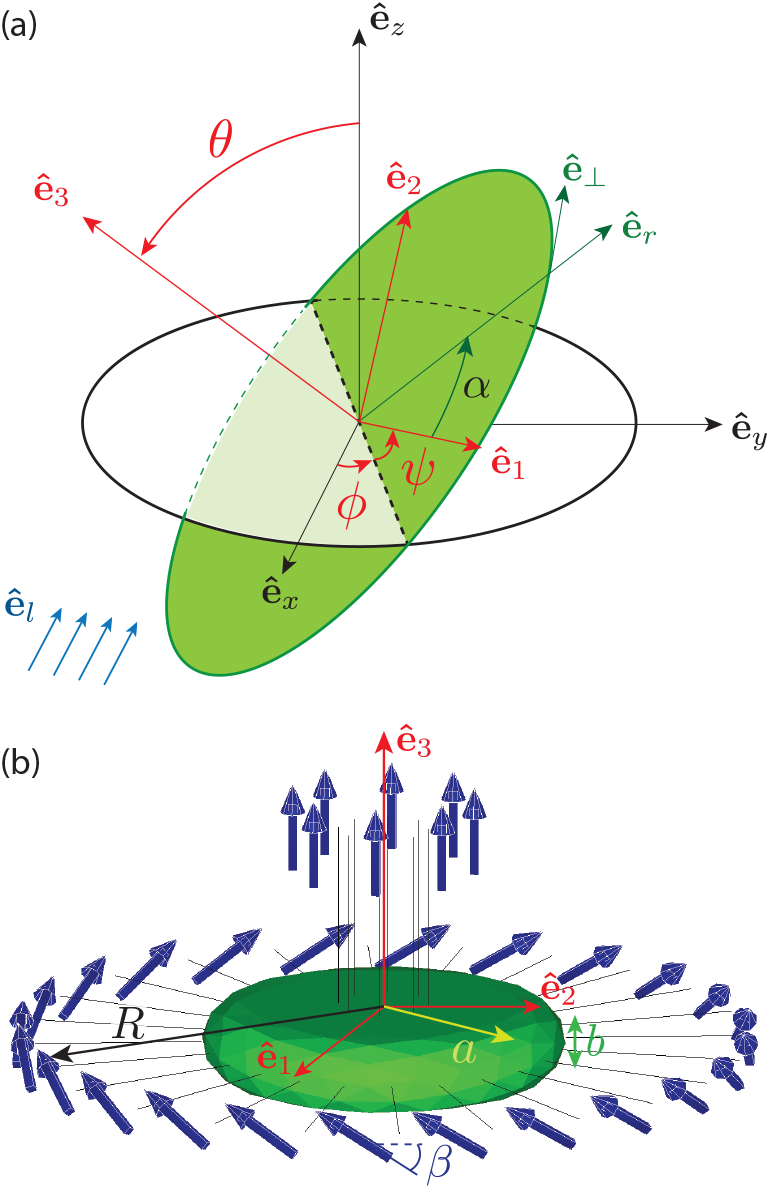
Details of computational geometry. (a) Coordinates system and Euler angles. The frame (**ê**_*x*_, **ê**_*y*_, **ê**_*z*_) is attached to the laboratory; the frame (**ê**_1_, **ê**_2_, **ê**_3_) is attached to the *Gonium* body (green disk), with **ê**_3_ the symmetry axis and (**ê**_1_, **ê**_2_) in the body plane. Euler angles (*θ, ϕ, ψ*) relate the two frames: by definition, *θ* is the angle between **ê**_*z*_ and **ê**_3_, *ϕ* is the angle from **ê**_*x*_ to the line of nodes (dotted line), and *ψ* is the angle from the line of nodes to **ê**_1_. In the *Gonium* body plane (**ê**_1_, **ê**_2_), the flagella are labeled by the angle *α*, with (**ê**_*r*_, **ê**_⊥_) the corresponding local frame such that cos *α* = **ê**_1_ · **ê**_*r*_. For the computation of the phototactic response, we assume the light is incident along **ê**_*l*_ = −**ê**_*x*_ (blue arrows). (b) *Gonium* geometry for simulations. The body (in green) is a thick disc with radius *a* = 20 *μ*m and thickness *b* = 8 *μ*m. Flagella (length 20 *μ*m) associated with a point force 20 *μ*m away from the cell body are attached to the body. The 8 central flagella generate thrust while the 24 peripheral ones are tilted by *β* ≃ 30°, and generate both thrust and rotation.

These viscous forces and torques are balanced by the thrust and spin induced by the action of the flagella. The central flagella produce a net thrust *F_c_* **ê**_3_ and no torque. The peripheral flagella contribute both to propulsion and rotation: we model them in the continuum limit as an angular density of force in the form *d***f**_*p*_ = **f**_*p*_*dα*, with **f**_*p*_(*α*) = *f*_*p*‖_ **ê**_3_ + *f*_*p*⊥_ **ê**_⊥_, where *α* is the angle labeling the flagella and **ê**_⊥_ = −sin *α***e**_1_ + cos *α***ê**_2_ the unit azimuthal vector along the *Gonium* periphery (Fig. 4a). Here, we choose *f*_*p*‖_ > 0 and *f*_*p*⊥_ < 0 to ensure a left-handed body rotation, with the ratio *f*_*p*‖_/*f*_*p*⊥_ = −tan *β* which we assume independent of *α*, and *β* the tilt angle made by the peripheral flagella (Fig. 4b). The flagella therefore produce a net force and a net torque

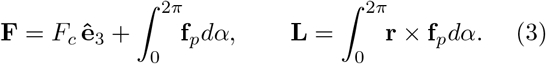

In the absence of phototactic cues and for a perfectly symmetric *Gonium*, **f**_*p*_ has no *α* dependence, and the force and torque are purely axial,

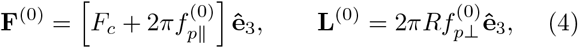

which satisfies **F**^(0)^ · **L**^(0)^ < 0. For such a *straight swimmer*, the velocity **U** and angular velocity **Ω** resulting from the balance with the frictional forces and torques are also purely axial, along **ê**_3_ and −**ê**_3_, respectively.

The helical trajectories observed experimentally indicate that in most *Gonium* colonies the forces **f**_*p*_(*α*) produced by the peripheral flagella are not perfectly balanced. Such an unbalanced distribution produces nonzero components of **F** and **L** normal to **ê**_3_, which deflects the swimming direction of the colony. The simplest imbalance compatible with the observed helical trajectories is a modulation of the force developed by the peripheral flagella in the form

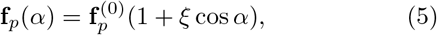

with 0 ≤ *ξ* ≤ 1 a parameter. With this choice, there is a stronger flagellar force along **ê**_1_ and the net force and torque acquire components in the plane (**ê**_2_, **ê**_3_). Interestingly, the combined effect of these transverse force and torque components is to enable such unbalanced colonies to maintain their overall swimming direction.

To relate the uneven distribution of forces (5) to the helical pitch angle *χ*, we consider the geometry sketched in Fig. 1d: a *Gonium* colony swims along a helix whose axis is along **ê**_*z*_, with **ê**_3_ describing a cone around **ê**_*z*_ of constant apex angle *ζ*. The angular velocity vector **Ω** is therefore along **ê**_*z*_. In terms of Euler angles (Fig. 4a), this choice implies *θ* = *ζ*, *ψ* = 0 and 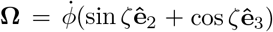 with constant 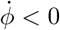 (See Appendix C).

The helix pitch angle *χ* is the angle between **U** and **ê**_*z*_. Because of the anisotropic resistance matrices and the axial force *F_c_* produced by the central flagella, *χ* is larger than the angle *ζ* between **ê**_3_ and **ê**_*z*_: the colony exhibits large lateral excursions while maintaining its symmetry axis **ê**_3_ nearly aligned with the mean swimming direction **ê**_*z*_. Expanding to first order in the imbalance amplitude *ξ*, we find from force and torque balance

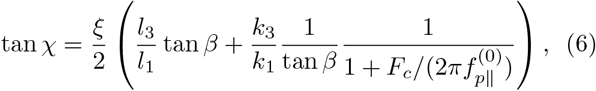

so a symmetric *Gonium* (*ξ* = 0) swims straight (*χ* = 0).

### C. Comparison with experiments and computations

The continuous angular density of forces considered in the previous section is convenient to obtain a simple analytical description of the helical trajectories. However, to estimate the main hydrodynamic properties (flagella force and resistance matrices) from measurements, a more realistic description is needed that considers the drag exerted on the cell body by the flow induced by the flagellar forces. We assume for simplicity that all flagella produce equal individual forces *F_i_*. With 4 central and 12 peripheral bi-flagellated cells, *F_c_* = 8*F_i_* for the central cells, and 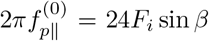 and 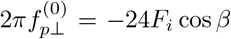 for the axial and azimuthal force developed by the peripheral cells. We first restrict our attention to *straight* swimmers with balanced peripheral flagella force (*ξ* = 0).

To identify the physical parameters (flagella force *F_i_*, flagella tilt angle *β*, effective radius *R*, translation and rotation friction coefficients *k*_1_, *k*_3_, *l*_1_, *l*_3_) from the experimental observations (swimming speed *v* ≃ 40 *μ*m/s and body rotation frequency *ν*_3_ ≃ 0.4 Hz), we combined computations and PIV measurements of the flow around a swimming *Gonium* colony. The computations, based on the boundary element method [33], are detailed in Appendix D. We use the simplified geometry of Fig. 4b: the cell body is a thick disk of radius *a* and thickness *b*, with 32 straight filaments representing the flagella pairs from the 16 cells, and a set of 32 point forces capturing the result of the flagella beating. These filaments also contribute to the hydrodynamic drag, and hence to the values of the friction coefficients *k*_1_, *k*_3_, *l*_1_, *l*_3_. Fagella lengths are 30 – 40 *μ*m, but we consider in the computations the average of their shape over a beat cycle as contributing to the drag, which we expect to be in the range of 20 – 30 *μ*m. Comparing the velocity fields from PIV and numerics (Figs. 5a,b), we identify the location of the point force at about 20 *μ*m from the cell body.

**FIG. 5.**
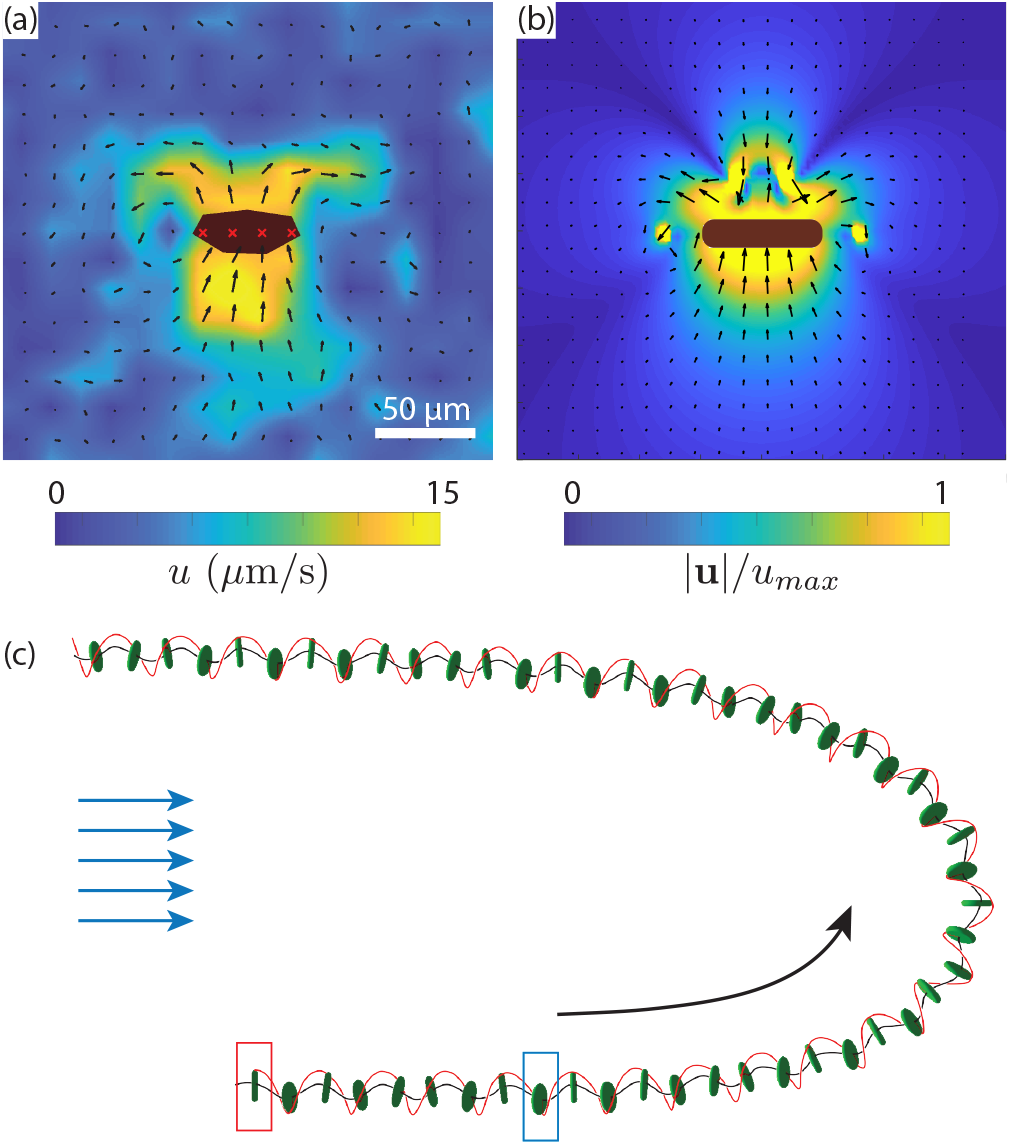
Photoresponse of *Gonium*. (a-b) Flow fields around a colony swimming towards the top of the image, in the laboratory frame. (a) Micro-PIV measurements (the body is in black with red crosses). The background color shows the norm of the velocity, varying here up to 15 *μ*m/s. Scale bar is 50 *μ*m. (b) Numerical flow field using the point-force model. (c) Wavy trajectory of a colony, numerically computed for *ξ* = 0.4 and *μs*_0_ = 0.5. The initial position is shown by the red rectangle, and the time when light is switched on from the left is indicated by the blue rectangle. Black line shows the trajectory, and the red line follows the position of the point of maximal force, *α* = 0, highlighting rotation of the body.

From this geometry, we compute the friction coefficients *k*_1_, *k*_3_, *l*_1_, *l*_3_ for several flagella lengths between 20 and 30 *μ*m, and tune the intensity of the point forces *F_i_* and tilt angle *β* to obtain a good match to the experimental swimming and angular speeds. Tuning the flagella length to 20 *μ*m, so that *R* ≃ 2*a* ≃ 40 *μ*m, appeared as the best match between experimental observations and numerical results. This corresponds to a point force per flagella *F_i_* ≃ 3.5 pN and a tilt angle *β* ≃ 30°. The friction coefficients are *k*_1_ ≃ 10, *k*_3_ ≃ 13, *l*_1_ ≃ 6, and *l*_3_ ≃ 8, as detailed in Appendix D. The flow field computed from these parameters (Fig. 5b) shows an overall structure in reasonable agreement with the PIV measurement performed around a freely swimming *Gonium* colony, which validates the methods. As expected, the far-field structure is typical of a *puller* swimmer, with inward flow in the swimming direction and outward flow normal to it.

Note that the computed force is the flagellar thrust only which does not take into account the drag force, so that it overestimates the total force exerted by a flagellum on the fluid. An estimate for the true force is obtained by balancing the viscous force experienced by a colony *F_v_* = −*k*_3_*ηRv* to the propulsive contributions of the flagella 8*F_i_* + 24*F_i_* sin *β* ≈ 20*F_i_*. The resulting force that a flagellum effectively applies on the fluid is of the order of 1 pN, a value close to the estimate for *Chlamydomonas* over a beating cycle [9].

We finally consider helical trajectories produced by unbalanced swimmers (*ξ* ≠ 0), and include in the computation angular modulation of the peripheral flagella force (5). A typical trajectory is illustrated in Fig. 5c and Supplementary Movie 3 (note the non-phototactic part of the trajectory, between the red and blue rectangles). The black and red lines show the helical body trajectory and the position of the maximal force (unit vector **ê**_1_). As expected, the anisotropic resistance matrices characteristic of the *Gonium* geometry produce a helical trajectory with large lateral excursion but moderate tilt of the body normal axis.

To relate the helical pitch angle *χ* to the parameter *ξ*, we extract from the simulated trajectories the helix amplitude *A*, wavelength λ and mean velocity *v_m_*, as described in the Appendix D. We deduce a pitch angle *χ*, observed to linearly increase with *ξ*, in close agreement with Eq. (6). The decrease in swimming efficiency with *χ* is also well reproduced, with numerical data closely following the geometrical prediction of *v_m_*/*v* = cos *χ*, as evidenced by the black dots in Fig. 3c. From the experimental measurement *χ* ≃ 30° ± 13°, we deduce an average amplitude *ξ* ≈ 0.4: the strongest flagella typically produce a force at least twice larger than the weakest ones. This surprisingly large value suggests that waviness is an important feature for the swimming of *Gonium*.

## III. PHOTOTACTIC SWIMMING

### A. Experimental observations

We now turn to the phototactic response of *Gonium*, which is triggered by adding to the previous experimental setup two blue LEDs on the sides of the chamber. The simplest configuration has the two lights arranged facing each other, as in Fig. 2a. Figure 6a displays the trajectories of a set of colonies reorienting as the two lights are alternately switched on and off for approximately 20s twice in a row, also seen in Supplementary Movie 4. These trajectories show various degrees of waviness, as in the non-phototactic experiments (Fig. 3a), however here their direction is no longer random but rather aligned with the incident light. At each change of light direction, a marked slowdown is also observed, as illustrated in Fig. 6b: just after the change in light, the swimming speed decreases by half for a few seconds, indicating a reduction in the flagella activity [25]. The variability in this drop can originate from out of plane swimming during reorientation, as we only have access to (*x, y*) projections of the trajectories.

**FIG. 6.**
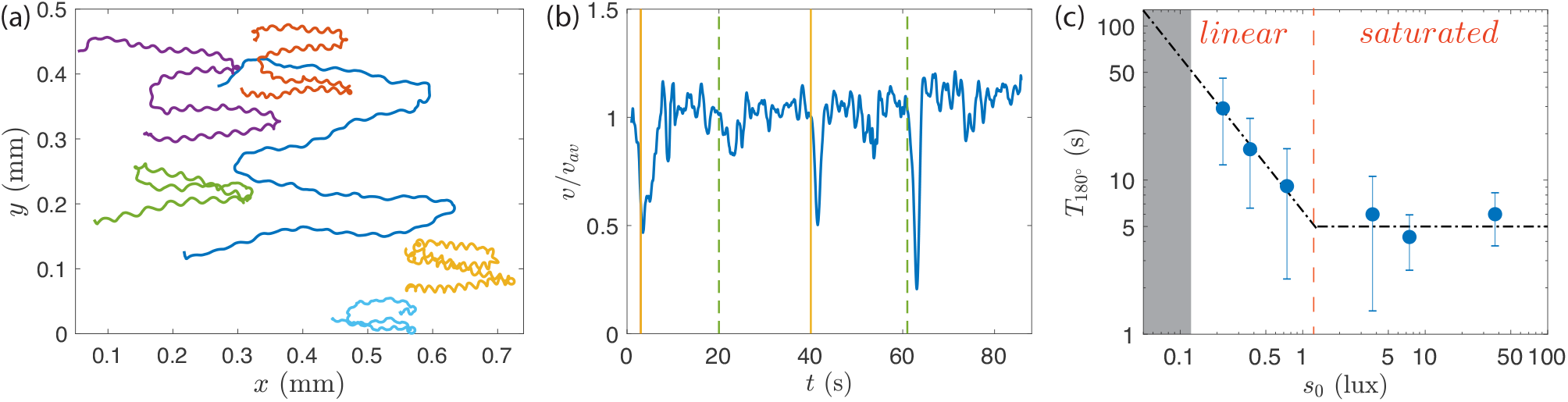
Phototactic reorientation of *Gonium*. (a) Reorientation trajectories of colonies under blue light stimulation shone alternatively from right to left twice during 20 s. Each color line corresponds to a single colony and shows a swimming direction alternating to the right and left according to the change in light source position. (b) Instantaneous normalized velocity as a function of time for the blue trajectory in (a). Vertical lines indicate a change in the light source position: yellow shows times when the light is shone from the right, while green dashed lines stand for light coming from the left. (c) Reorientation time *T*_180°_ as a function of the light intensity *s*_0_. The line shows *T*_180°_ ≃ 1/*s*_0_ for so < 1 lux, consistent with Eq. (15), and a saturation at *T*_180°_ = 5 s at larger *s*_0_. The grey shaded area corresponds to times longer than the video-camera trigger.

A key feature to model the dynamics of the body reorientation towards light is the dependence of the flagella force on light intensity. We measured the time *T*_180°_ to perform a turn-over as a function of the light intensity *s*_0_ (Fig. 6c), restricting measurements to *s*_0_ < 100 lux, so as to observe only *positive* phototaxis; at larger *s*_0_, an increasing fraction of colonies display *negative* phototaxis, swimming away from the light (the critical light intensity between positive/negative phototaxis varies with time during the diurnal cycle).

Figure 6c shows two phototactic regimes: A *linear*regime at moderate intensity (*s*_0_ < 1 lux), for which *T*_180°_ ~ 1/*s*_0_, and a *saturated* regime at larger *s*_0_, for which *T*_180°_ ≃ 5 s ± 1 s. In what follows we focus on the linear regime, in which the flagella activity is proportional to the light intensity. The constant reorientation time in the saturated regime may originate from a biological saturation in the signal transmission from the eye-spot to the flagella, or from a hydrodynamic limitation in the reorientation process itself: In this saturation regime, we have *ν*_3_*T*_180°_ ≃ 2, which is probably the fastest reorientation possible – during a 90° reorientation the *Gonium* colony performs one single rotation, *i.e*. each eye-spot detects the light variations only once.

A direct consequence of this dependence on light intensity is the shape of the trajectories during the reorientation process, illustrated in Fig. 7; here, only the reorienting part of the trajectories (*t* > 0 s) is displayed, and all are centered at (0,0) when the light is switched on from the left. At *s*_0_ = 0.4 lux, colonies swim a long distance before facing the light, whereas they turn much more sharply at larger *s*_0_. The similar trajectories observed for *s*_0_ = 4 lux and 40 lux are consistent with saturation in phototactic response for *s*_0_ > 1 lux seen in Fig. 6c. Note that at *s*_0_ = 40 lux, a small fraction of colonies (typically 10%) start to show erratic trajectories with a weak negative phototactic component.

**FIG. 7.**
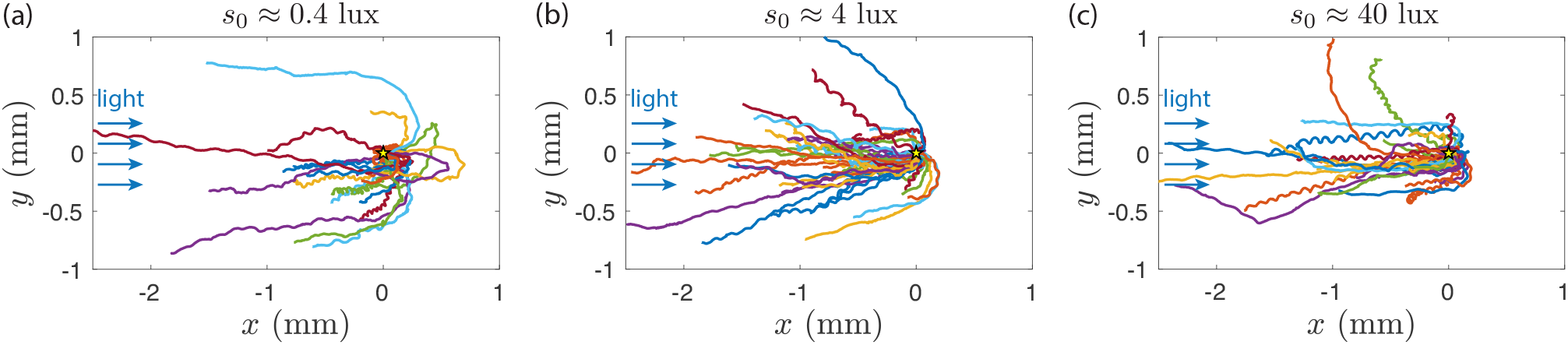
Phototactic turn-over after a change of light incidence. *Gonium* colonies are initially swimming (*t* < 0 s) toward a light of constant intensity on the right. At *t* = 0 s, this light is switched off while another of controlled adjustable intensity *s*_0_ is shone from the left. Trajectories are reported for *t* > 0 s, and have been shifted to the origin at *t* = 0 s.

A remarkable feature of Fig. 7 is that wavy trajectories gather near the center line, indicating a quick change in orientation, while smoother trajectories show a larger radius of curvature. This suggests that waviness, detrimental for swimming efficiency, is beneficial for phototactic efficiency: by providing a better scan of their environement, eye-spots from strongly unbalanced *Gonium* may better detect the light variations.

### B. Reaction to a step-up in light

To explain the reorientation trajectories, we need a description of the flagella response to time-dependent variation in the light intensity. Measuring this response in freely swimming colonies is not possible because of their complex three-dimensional trajectories. To circumvent this difficulty, we use as in previous work [7, 13], a micropipette technique to maintain a steady view of the colony. Because of the linearity of the Stokes flow, the fluid velocity induced by the flagellar action is proportional to the force they exert.

The experimental setup is described in Fig. 2b and Appendix A. Micropipettes of inner diameter slightly smaller than the *Gonium* body size (Fig. 8a) are used to catch colonies by gentle aspiration of fluid. An optical fiber connected to a blue LED is introduced in the chamber, and micro-PIV measurements are performed to quantify the changes in velocity field around the colony while varying the light intensity. Due to the presence of the micro-pipette, the measured velocities are not reliable on the pipette side (right-hand side of the images), but are accurate in the remainder of the field of view.

**FIG. 8.**
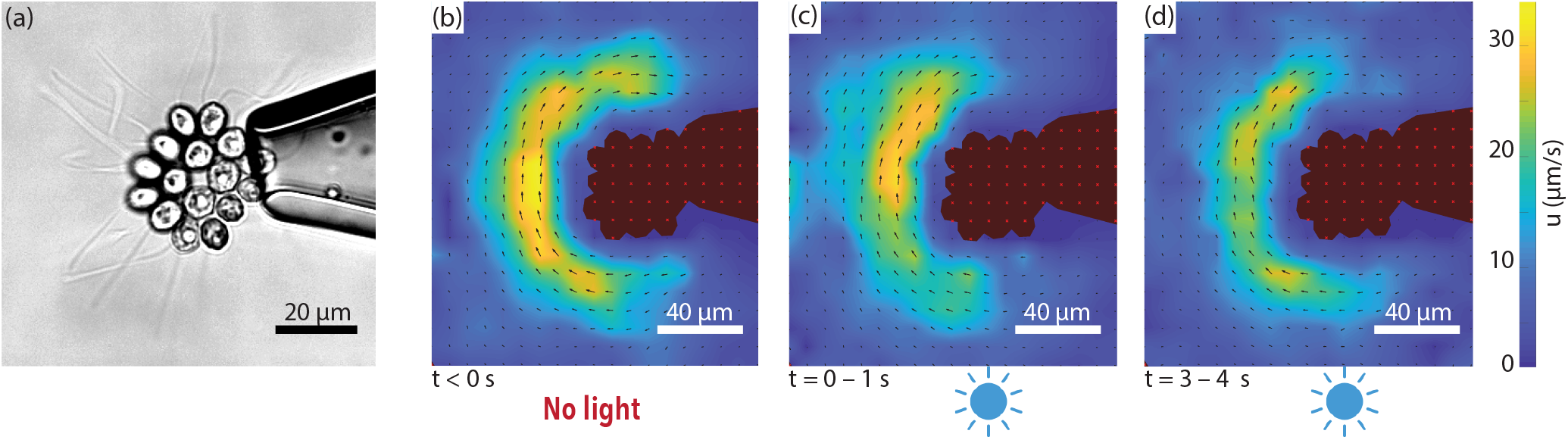
Micropipette experiments. (a) A 16-cell colony held on a micro-pipette, viewed from posterior side. Flagella are clearly visible and their frequency can be followed as a function of time and light. The blue LED is located in the same plane as the micropipette, a few millimeters below the bottom of the image. (b) - (d) Velocity fields measured by micro-PIV, averaged over 1 s at three different times indicated below each panel. The *Gonium* is seen from the back (flagella away from us). Colormap is the same across the three images, from 0 – 30 *μ*m/s. Light intensity *s*_0_ ≈ 1 lux.

In these experiments we focus on phototactic stimulation in the form of a step-up in intensity (see also Supplementary Movie 5). The elementary flagella response to this is useful to compute the response to a more realistic change experienced by the flagella during the reorientation process. The micro-PIV experiments in Fig. 8b-d show the flow around a *Gonium* at three key moments of a step-up experiment: prior to light excitation, the flow is nearly homogeneous and circular, with peak velocities of 30 *μ*m/s at a distance of 20 – 30 *μ*m from the cell body. Immediately after light is shone from the bottom of the image (*i.e*. at 90° to the *Gonium* body plane), the flow symmetry is clearly broken: the velocity is strongly reduced in the illuminated part, down to ≃ 15 *μ*m/s, about half the original value, while it remains essentially unchanged in the shadowed part. Finally, after a few seconds of constant illumination, the velocity gradually increases and the initial symmetry of the flow field is eventually recovered: the phototactic response is adaptive to the new light environment.

Flow velocities measured on the illuminated and shadowed sides are compared in Fig. 9a. Before light is switched on at *t* = 0, both curves follow similar variations around 30 *μ*m/s. At *t* = 0 s, the velocity on the illuminated side drops, while that on the opposite side remains nearly constant. After a few seconds, they tend to merge and variations are closer. As the flow velocity is proportional to the force in Stokes regime, this demonstrates a rapid drop followed by a slow recovery in the force from the illuminated side, while the force in the shadowed side remains essentially unaltered. We note that these PIV measurements only provide information on the *azimuthal* component of the force *f*_*p*⊥_, whereas the phototactic torque (3) is related to a non-axisymmetric distribution of the *axial* component of the force *f*_*p*‖_. However, because of the small tilt angle of the flagella, *β* ≃ 30°, changes in *f*_*p*‖_ are too difficult to detect experimentally by PIV. We assume here that the angle *β* is not impacted by light, so that measurements of the azimuthal flow variations provide a good proxy for the axial flow variations.

**FIG. 9.**
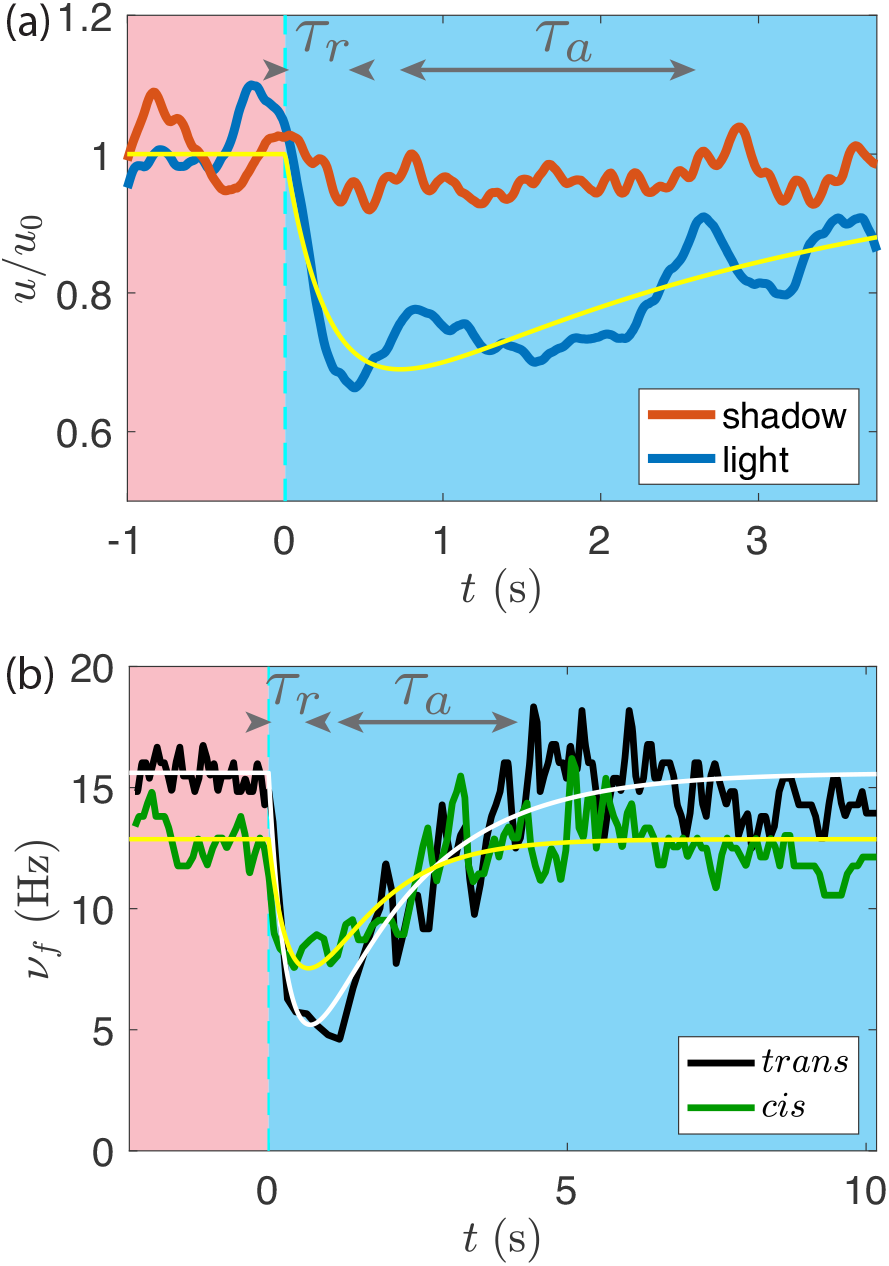
Adaptive phototactic response. (a) Normalized azimuthal velocities in the shaded (red curve) and illuminated (blue curve) sides of a colony held on a micropipette (averages over two similar experiments). For *t* < 0 s there is only red illumination (no phototaxis). Blue light is switched on at *t* = 0 s. The yellow line displays the adaptive model, Eq. (7) and (9). (b) *Trans* and *cis* flagella beat frequencies *ν_f_* (in black and green respectively), of a single cell receiving light (at *s*_0_ ≈ 1 lux) as a function of time. White and yellow lines are the respective best fits with Eqs. (7) and (9).

The velocity induced by a flagellum (hence the applied force) is a complex combination of beat frequency and waveform [34]. These two quantities can be measured only in simple geometries, such as in *Chlamydomonas*, where the two flagella beat in the same plane. The complex three-dimensional flagellar organization in *Gonium*makes it difficult to quantify changes in waveform, but the response in beat frequency of each individual flagellum can be readily measured. A typical response to a step-up is displayed in Fig. 9b for the two flagella (*cis* and *trans*) of a single cell detecting the light. The typical drop-and-recovery pattern of the velocity response is also remarkably present in the beating frequency, showing that the force induced by the flagella is governed at least in part by the beat frequency. The initial frequency without light stimulation is about 15 Hz, with slightly lower values systematically found for the *cis*-flagellum (close to the eye-spot; see Fig. 1b). Both flagella show a reduced beating frequency when light is switched on, with a more pronounced drop for the *trans*-flagellum. In *Chlamydomonas*, the *cis*-flagellum shows a strong *decrease* [16, 30] while the *trans*-flagellum slightly *increases* its frequency, a behavior which we do not observe in *Gonium*. Although a *cis - trans* differentiation is the key to phototaxis in *Chlamydomonas*, this trait, not required for phototaxis at the level of the colony, is also present at the cell level in *Gonium*.

### C. Adaptive model

Here we relate the drop-and-recovery response of the illuminated flagella to the phototactic reorientation. Whereas in earlier work on *Volvox* phototaxis [7] we considered the direct effect of changes in flagellar beating on the local fluid *velocity* on the surface, adopting a perpsective very much like that in Lighthill’s squirmer model [35], here we model the effect of light stimulation on the *force* developed by a peripheral flagella at an angle *α* as

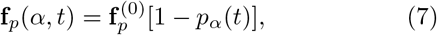

with *p_α_* the phototactic response. We neglect the possible phototactic response of the central flagella, which presumably do not contribute to the reorientation torque. We first describe the time-dependence of the response of a given flagellum *α* experiencing a step-up in light intensity, and then integrate the response from all flagella, taking into account eye-spot rotation, to deduce the reorientational torque, along the lines in recent work on *Chlamydomonas* [13].

The drop-and-recovery flagella response suggests using an adaptive model, which has found application in the description of sperm chemotaxis [36] and in phototaxis of both *Volvox* [7] and *Chlamydomonas* [13]: we assume that *p*(*t*) follows the light stimulation on a rapid timescale *τ_r_*, and is inhibited by an internal chemical process on a slower timescale *τ_a_* described by a hidden variable *h*(*t*). The phototactic response is assumed proportional to the light intensity, so we restrict ourselves to the linear regime at low *s*_0_. The two quantities *h* and *p* obey a set of coupled ODEs,

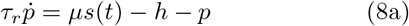

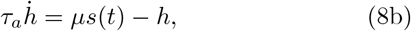

where *μ* is a factor with units reciprocal to those of *s*(*t*). This factor represents the biological processes which link the detection of light to the subsequent physical response. In the case of a step-up in light stimulation, *s*(*t*) = *s*_0_ *H*(*t*), with *H* the Heaviside function, we have

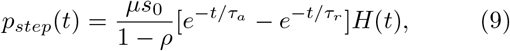

with *ρ* = *τ_r_*/*τ_a_*. This phototactic response (7)-(9), plotted as thin lines in Fig. 9 in the case *s*_0_ ≈ 1 lux, provides a reasonable description of the velocity and beating frequency, with *τ_r_* ≃ 0.4 ± 0.1 s and *τ_a_* ≃ 1.5 ± 0.5 s.

From this fit, we infer the value of the phototactic response factor: *μ* ≃ 0.6 and 0.8±0.1 lux^−1^ for the *cis* and *trans* flagella respectively. A somewhat lower value is obtained from the velocity signal, *μ* ≃ 0.4 ± 0.1 lux^−1^, which probably results from an average over the set of flagella on the illuminated side and a possible influence of a change in the flagella beating waveform. This disparity in the evaluation of *μ* underlines the complexity of the biological processes this variable summarizes. We retain in the following an average value *μ* ≃ 0.6 ± 0.2 lux^−1^.

By linearity, the phototactic response to an arbitrary light stimulation *s*(*t*) can be obtained as a convolution of the step-up response,

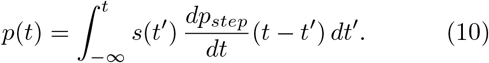

During the reorientation process, the light *s*(*t*) perceived by the peripheral cells varies on a time scale 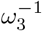. According to Eq. (10), the response to this light variation is band-pass filtered between *τ_a_* and *τ_r_*: an efficient response (*i.e*., a short colony reorientation time) is naturally expected for 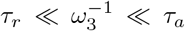. As shown in Appendix E, the maximum amplitude of the phototactic response is found in the limit *τ_r_*/*τ_a_* → 0 in Eq. (9), yielding *p*(*t*) = *μs*(*t*), which would correspond to a response following precisely the light stimulation. From our data, shown in Fig. 10, this behaviour has apparently not been selected by evolution, and we observe large fluctuations in the characteristic times *τ_r_* and *τ_a_*, suggesting other evolutionary advantages linked to this behaviour. Note that in *Chlamydomonas*, whose eye-spot is located at 45° from the *cis*–flagellum [37], an additional delay *τ_d_* associated with body rotation is needed between the light detection by the eye-spot and activation of the flagellum.

**FIG. 10.**
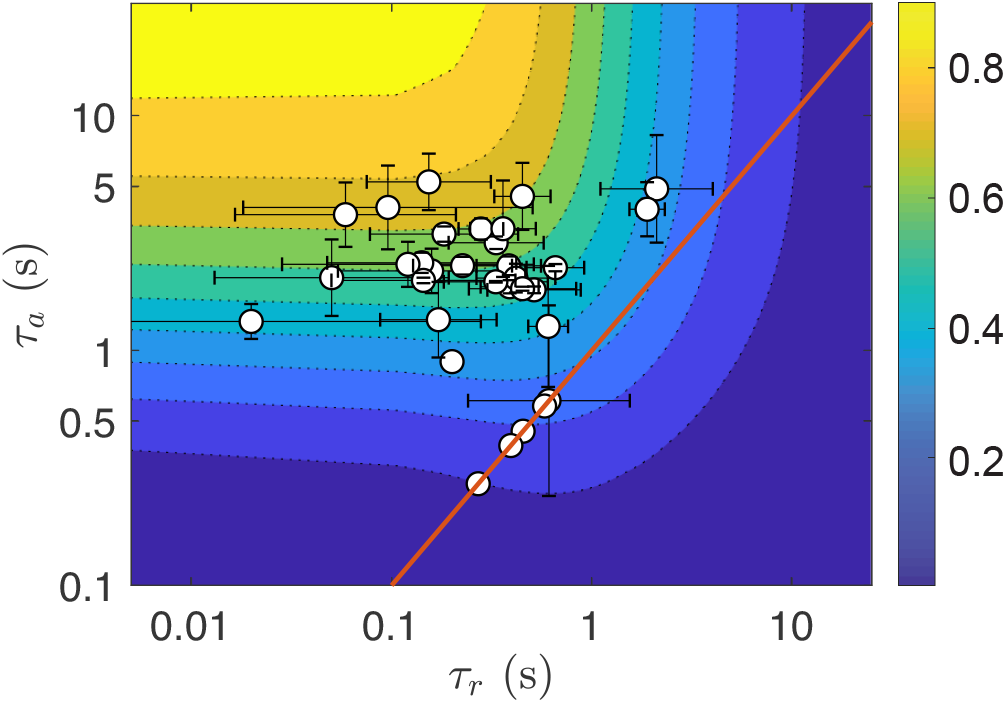
Experimental characteristic times (*τ_r_, τ_a_*) of the adaptive model. Data obtained from the flagella beat frequencies averaged for each colony (sample size: 34 colonies). Background color shows the gain function 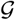 defined in Appendix F, Eq. (F8). Red line indicates the relation *τ_r_* = *τ_a_*.

To compute reorienting trajectories we consider for simplicity a perfectly balanced colony, with *ξ* = 0. In terms of phototactic reorientation, this situation is somewhat singular: such a hypothetical colony would swim in straight line, so that a change of 180° in the light incidence, precisely opposed to their swimming direction, could not be detected by the peripheral eyespots. We therefore model a reorientation with a light incidence **ê**_*l*_ at 90° to the initial swimming direction; this is the optimal configuration for light detection. We consider in the following **ê**_*l*_ = −**ê**_*x*_, and an initial orientation **ê**_3_ = −**ê**_*y*_, yielding *θ* = *π*/2 and *ϕ* = 0 (see Fig. 4). We assume for simplicity that *θ* remains constant during the reorientation: the colony axis **ê**_3_ rotates only in the plane (**ê**_*x*_, **ê**_*y*_) and the phototactic torque is along **ê**_*z*_ only. At the end of the reorientation process, we have **ê**_3_ = **ê**_*x*_, as the *Gonium* is then facing the light, *i.e. ϕ* = *π*/2.

The light perceived by a cell is

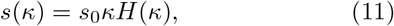

with *κ* = −**ê**_*ℓ*_ · **ê**_*r*_ the projected light incidence along the local normal unit vector **ê**_*r*_,

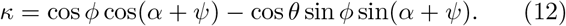

From Eq. (10), at a given time *t*, each flagellum labeled by the angle *α* has a phototactic response *p_α_*(*t*) resulting from the light *s*(*t*′) perceived at all times *t*′ < *t*. This light intensity depends on its orientation, as described by Eq. (11)–(12): it originates both from the (fast) body spin 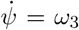 and the (slow) reorientation angular velocity 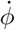. For simplicity, the computation of *p_α_* (given in Appendix F) assumes 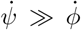, which is a valid approximation in the linear regime (*s*_0_ < 1 lux). Using the force (7), the reorientational torque is

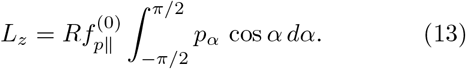

Balancing this torque with the vertical component of the frictional torque, 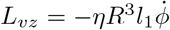, leads to the o.d.e

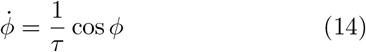

with a relaxation time expressed as a product of three factors,

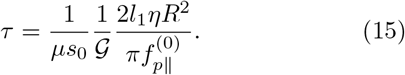

Here, *τ* ~ 1/*μs*_0_ is a dependence consistent with the linear response assumption, and we have introduced a gain function 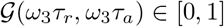 associated with the adaptive response, which is function of the non-dimensional relaxation times *ω*_3_*τ_a_* and *ω*_3_*τ_r_*, and described in Appendix F. This gain function is such that 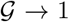 in the optimal case *ω*_3_*τ_r_* → 0 and *ω*_3_*τ_a_* → ∞ (no adaptive filtering), yielding the fastest reorientation time. With the pN scale of forces, the ~ 20 – 30*μ*m scale of *R* and the viscosity of water, we naturally find a timescale on the order of seconds, as in experiment.

Equation (14) is analogous to the problem of a door pulled by a constant force with viscous friction. Its solution with initial condition *ϕ*(0) = 0 is

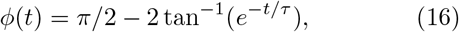

which asymptotes to *ϕ*(∞) = *π*/2: we have obtained the expected phototactic reorientation, on a timescale *τ*. In the case of a full reorientation (180°), this solution applies only for the second half of the reorientation, for *ϕ*(*t*) increasing from 0 to *π*/2 (the first half is simply deduced by symmetrizing Eq. (16) for *t* < 0).

### D. Comparison to the experiments

To study how far *Gonium* colonies are from the optimum gain 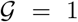, we plot in Fig. 10 the adaptive timescales (*τ_r_, τ_a_*) measured for a set of colonies under illumination intensity *s*_0_ ≈ 1 lux. These data were obtained by averaging the response times of the beating frequency of each flagellum. The measurements are centered around (*τ_r_, τ_a_*) ≈ (0.1, 2) s, corresponding to values of 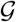 between 0. 5 and 0. 7, which is in the upper half, but somewhat far from the optimum expected for an ideal swimmer. The error bars highlight the variability in the response times among flagella within a colony, and this evidences how biological variations divert *Gonium* from its physical optimum.

Until now, we have ignored the helical nature of the trajectories in the phototactic reorientation process. Including waviness in the adaptive model of Sec. IIIC would be a considerable analytical task, because the light variation perceived by each eyespot would depend on the three Euler angles and their time derivatives. Instead, we performed a series of direct numerical studies of phototactic reorientation using the computational model previously introduced, combining in the flagella force the azimuthal modulation (5) and the phototactic response (7). Typical trajectories, obtained for various imbalance parameter *ξ* (and hence pitch angle *χ*), are illustrated in Fig. 11a and Supplementary Movie 3, for a light intensity corresponding to *μs*_0_ = 0.5. The trajectories clearly show a faster reorientation for wavier colonies, consistent with the experimental observations in Fig. 7. The trajectories show about 10 body rotations during the reorientation (i.e., 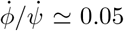), indicating that the linear regime assumption used in the model is satisfied for this value of *μs*_0_.

**FIG. 11.**
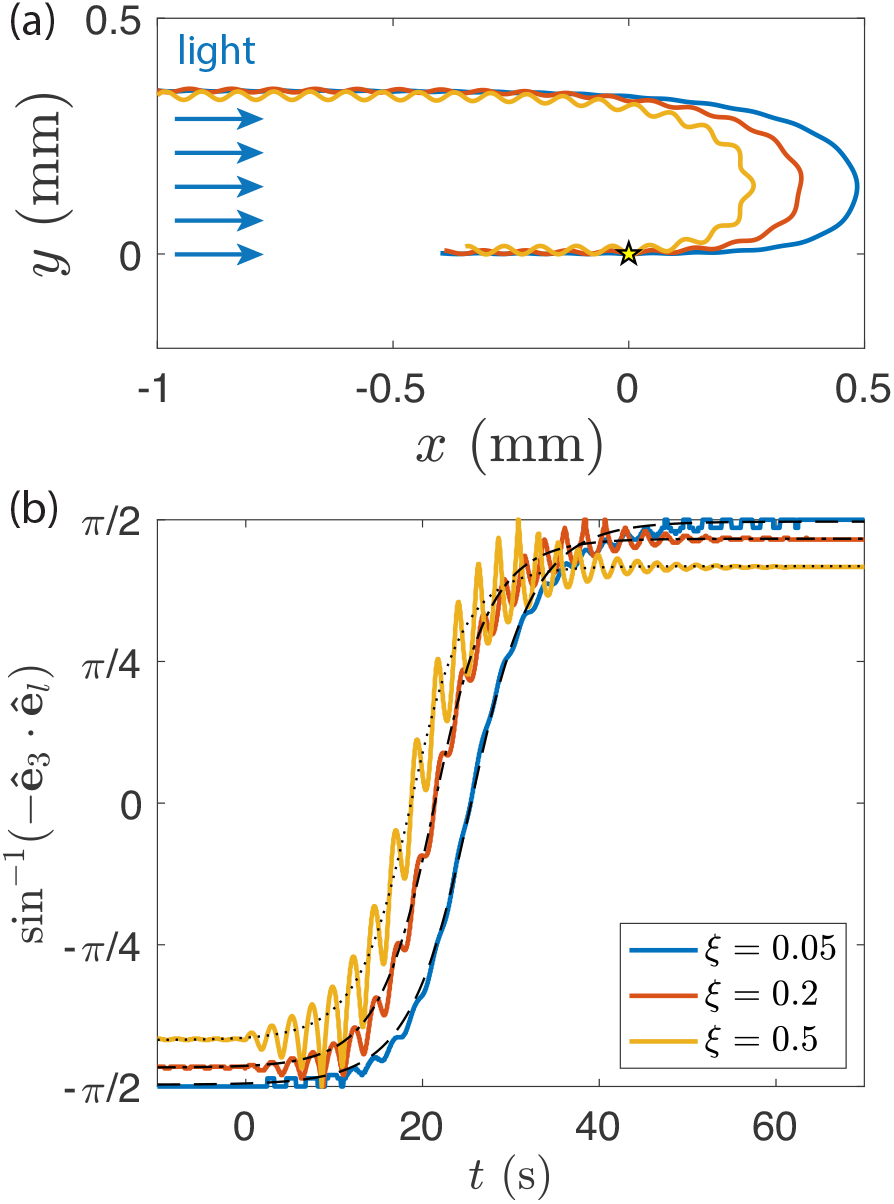
Numerical results for the phototactic response. (a) The reorientation trajectories of three wavy swimmers are shown for *μs*_0_ = 0.5 (corresponding to *s*_0_ ≈ 0.3 lux). They are initially swimming away from the light source, and then turn around after the light is switched on from the left when colonies are at the origin (yellow star). (b) Evolution of *ϕ* as a function of time for the same swimmers. Light is turned on at *t* = 0. The thin black dashed, dash-dotted and dotted lines are the respective fits using Eq. (16).

The reorientation dynamics of such wavy swimmers is illustrated in Fig. 11b, showing the angle sin^−1^(−**ê**_3_ · **ê**_*l*_) as a function of time. For a straight swimmer, this angle is simply *ϕ*(*t*): it increases monotonically from −*π*/2 to *π*/2 following Eq. (16) symmetrized in time. For an unbalanced swimmer, the symmetry axis **ê**_3_ describes a cone of apex angle *ζ* (as described in Sec. II) around the mean swimming direction: sin^−1^(−**ê**_3_ · **ê**_*l*_) therefore increases from −*π*/2+*ζ* to *π*/2−*ζ*. The angle obtained from numerical simulations indeed shows oscillations superimposed on a mean evolution that is still remarkably described by the straight-swimmer law [Eq. (16), in dotted lines], with the asymptotic values ±*π*/2 simply replaced by ±(*π*/2 − *ζ*). Increasing the imbalance parameter *ξ* clearly increases the amplitude of the oscillations when the light is switched on, yielding a better scan of the environment and hence a shorter reorientation timescale *τ*. Combined with the reduced mean velocity *v_m_* = *v*/ cos *χ* (Fig. 3c), this faster reorientation finally produces the sharper trajectories observed in Fig. 11a. Our simulations therefore successfully capture the main features of the phototactic reorientation dynamics observed in *Gonium* colonies.

## IV. CONCLUSIONS

We have presented a detailed study of motility and phototaxis of *Gonium pectorale*, an organism of intermediate complexity within the *Volvocine* green algae. In its flagellar dynamics it combines beating patterns found in *Chlamydomonas* and *Volvox*, and has a distinct symmetry compared to the approximate bilateral symmetry of the former and the axisymmetry of the latter. Our experimental observations are consistent with a theory based on adaptive response exhibited solely by the peripheral cells, on a time scale comparable to the rotation period of the colony around the axis normal to the body plane. The precise biochemical pathways that underlie this adaptive response remain unclear. As with other green algae [7, 13], the response and adaptation dynamics serve to define a kind of bandpass filter of response centered around the colony rotation period, extending from tenths of a second to several seconds. It is natural to imagine that evolution has chosen these scales to filter out environmental fluctuations that are both very rapid (such as might occur from undulations of the water’s surface above a colony) and very slow (say, due to passing clouds), to yield distraction-free phototaxis. Finally, we have observed that colonies with helical trajectories that arise from slightly imbalanced flagellar forces are shown to have enhanced reorientation dynamics relative to perfectly symmetric colonies. Taken together, these experimental and theoretical observations lend support to a growing body of evidence [38] suggesting that helical swimming by tactic organisms is not only common in Nature, but possesses intrinsic biological advantages.

## ACKNOWLEDGMENTS

We thank Kyriacos Leptos for initial experimental assistance and inspiration for the model, Lucie Domino for preliminary studies of *Gonium*, and David Page-Croft, Caroline Kemp, and John Milton for vital technical support. We are also grateful to Matt Herron and Shota Yamashita for insights into *Gonium* structure. This work was supported in part by Wellcome Trust Investigator Award 207510/Z/17/Z (REG & HdM), Institut Universitaire de France (FM), the Japan Society for the Promotion of Science (KAKENHI Grant Nos. 17H00853 & 17KK0080 to TI), Established Career Fellowship EP/M017982/1 from the Engineering and Physical Sciences Research Council and Grant No. 7523 from the Gordon and Betty Moore Foundation (REG).

## Appendix A: Materials and methods

### Culture conditions

Wild-type *Gonium pectorale* colonies (strain CCAC 3275 B from the Cologne Biocenter) were grown in standard *Volvox* medium in an incubator (Binder) at 24°, under 3800 lux illumination in a 14:10h light-dark cycle.

### Observation of swimming

Chambers were prepared using two glass slides, sealed with Frame-Seal Incubation Chambers (SLF0201, 9×9 mm, BIO-RAD Laboratories). They were subsequently mounted on a Nikon TE2000-U inverted microscope for observation with a 4× or 10× Nikon Plan Fluor objective. Non-phototactic red illumination of the samples was achieved with a long pass filter (620 nm) in the light path. Observations were recorded with a high-speed video camera (Phantom V311; see also below) at 30 fps. A pair of blue LEDs (M470L2, Thorlabs), connected to LED drivers (LEDD1B, Thorlabs), were placed on either sides of the chamber, and separately controlled, as described in Fig. 2a. To assure maximum photoresponse, the wavelength of 470 nm was chosen to be close to the absorption maximum rhodopsin [39], the protein in the eyespot responsible for light detection [14].

### Micropipette experiments

We used borosilicate glass pipettes (Sutter Instruments, outer diameter 1.0 mm, inner diameter 0.75 mm), a pipette puller (P-97, Sutter Instruments) and a multifunction microforge controller (DMF1000, World Precision Instruments) to produce micropipettes of inner diameter ~ 20 *μ*m, with polished edges.

Chambers were fabricated by gluing short spacers to glass slides with UV-setting glue (NOA61), with UV exposure for 1 min (ELC-500 lamp, Electro-Lit Corporation), leaving apertures at 90° for the micropipette and the optical fiber to enter the chamber, as sketched in Fig. 2b. Colonies were caught on the micropipette, through which fluid was gently aspirated via a 10 ml syringe (BD Luer-Lok 305959). The optical fiber (FT400EMT, Thorlabs) wass connected to a 470 nm LED (M470F3, Thorlabs) and driver (DC2200, Thorlabs), enabling intensity and timing control. We took care that the fiber end enters the liquid in the chamber to avoid losses of light by reflection at the air-water interface, and that it is aligned with the tip of the micropipette. Synchronization of the light source to the camera wass made via a NI-DAQ (BNC-2110, National Instruments). Light intensities were measured with a lux meter (Lutron LX-101). We verified proportionality between the intensity (in lux) from of the optical fiber and the current (mA) provided by the driver, to extrapolate light intensities below 1 lux, where the meter saturates.

Microparticle image velocimetry (PIV) experiments are conducted by adding non-fluorescent beads (Polybead Polystyrene Cat. 07310, Polysciences), of diameter 1 *μ*m, to the *Gonium* suspension. Image acquisition was performed on a similar microscope with a 20 × Plan Fluor (Nikon) or a 63× water immersion Plan Apochromat objective (Zeiss) with a 45 – 60 Nikon adapter, connected to the same video-camera recording at 200 fps. Image analysis was performed via the Matlab tool PIVlab, with a window size adapted to the chosen microscope objective to ensure the presence of at least 3 – 5 particles.

## Appendix B: Inferring Helical Motion from Planar Projections

We define three velocities for a helical trajectory: the mean velocity *v_m_* = λ/*T*, where *T* is the time to cover one wavelength λ, the true (3D) instantaneous velocity *v*_3 *D*_ and the apparent (2D) instantaneous velocity *v*_2 *D*_ = *v*. From microscopy, we access *v_m_* and *v*, from which we deduce *v*_3 *D*_. To relate these velocities, we rewrite the helical trajectory in parametric form:

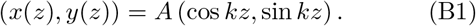

where *k* = 2*π*/λ. The instantaneous velocity is *v*_3 *D*_ = *l*/*T*, where 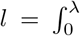 is the curved length covered by *Gonium*. In 3D, we have 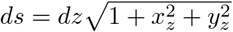, yielding

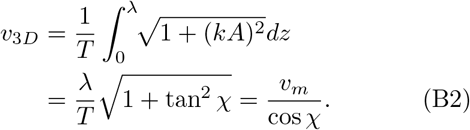

If we assume that the 2D plane of view is (*x, z*), then for the apparent (2D) instantaneous velocity *v* we now have 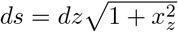, so

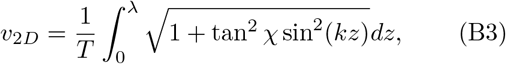

which is an elliptic integral of the second kind of parameter tan^2^ *χ*. For small *χ*, a Taylor expansion gives *v_m_*/*v*_3 *D*_ ≃ 1 − *χ*^2^/2 and *v_m_*/*v*_2 *D*_ ≃ 1 − *χ*^2^/4. For the typical values found for *Gonium, χ* ≈ 30°, we deduce that *v*_2 *D*_/*v*_3 *D*_ ≃ 0.93, and therefore conclude that approximating the instantaneous velocity by its 2D projection is reasonable.

## Appendix C: Helical trajectories

We relate here the pitch angle *χ* of the helical trajectories to the uneven distribution of flagellar forces. The geometry is shown in Fig. 1d, with **ê**_3_ the symmetry axis of the *Gonium* body. In this geometry, the rotation vector **Ω** is along the mean swimming direction **ê**_*z*_. The pitch angle *χ* is the angle between the instantaneous velocity **U** and **ê**_*z*_. The symmetry axis of the *Gonium* body describes a cone around *z* of constant apex angle *ζ*, with *ζ* < *χ* because of the anisotropic resistance matrices. The Euler angles for this geometry are *θ* = *ζ*, 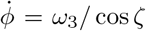 and 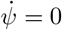. The frictional torque (1b) is thus

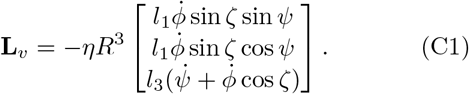

The simplest defect compatible with such a helical trajectory is a modulation of the axial force developed by the peripheral flagella in the form (c.f. Eq. (5))

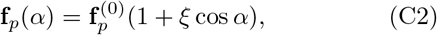

with *ξ* a “defect” parameter. We assume here that *β* does not depend on *α*, so that the defect affects the peripheral and axial components *f*_*p*‖_ and *f*_*p*⊥_ in the same way. From this force distribution, the total force and torque (3) are

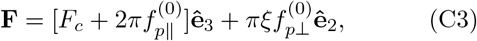

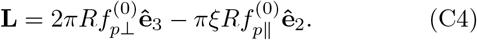

Since the force and the torque now have components along **ê**_2_, the velocity and angular velocity do as well. The phase choice in Eq. (5) assigns maximum flagella force along **ê**_1_, and implies *L*_1_ = 0, so *ψ* = 0 in Eq. (C1).

To compute the angle *ζ*, we first solve for the torque-angular velocity relation along **ê**_2_ and **ê**_3_,

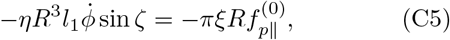

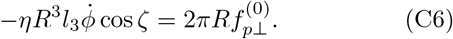

Solving for *ζ* yields

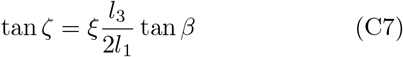

with 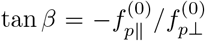. A symmetric *Gonium* (*ξ* = 0) naturally has *ζ* = 0.

The helix pitch angle *χ*, the angle between the instantaneous velocity **U** and the mean swimming direction **ê**_*z*_, is found by solving the velocity-force relation along **ê**_2_ and **ê**_3_,

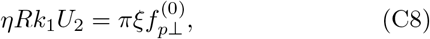

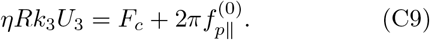

The angle *ζ*′ between **ê**_3_ and **U** is

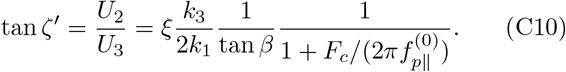

We note that the vectors **Ω** (‖ **ê**_*z*_), **ê**_3_ and **U** are in the same plane (**ê**_2_, **ê**_3_). The total angle *χ* between **ê**_*z*_ and **U** is therefore simply *χ* = *ζ* + *ζ*′. For small *ξ*, we have tan *χ* ≃ tan *ζ* + tan *ζ*′, yielding Eq. (6),

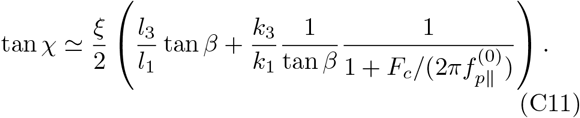

We can rewrite this relation using the discret point-force model. Assuming that all the individual flagellar forces *F_i_* are identical, we have *F_c_* = 8*F_i_* and 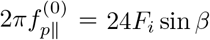, yielding

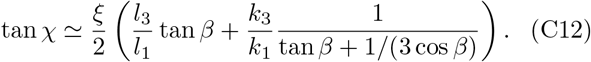

The helix parameters *A* and λ are derived by expressing the velocity in the laboratory frame,

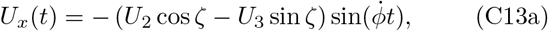

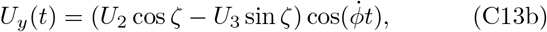

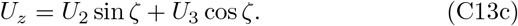

with 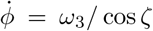. Here, *U_z_* corresponds to *v_m_*, the mean velocity along *z*. Integrating these equations in time yields the helix amplitude and wavelength,

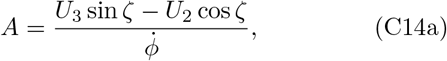

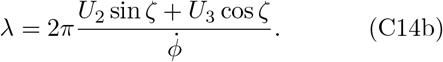

Using tan *χ* = 2*πA*/λ, we obtain

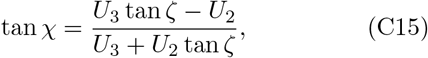

which can be shown to be equivalent to Eq. (C12) to first order in *ξ*.

## Appendix D: Numerical methods

Here we give details of the numerical techniques and show characteristic values extracted from the computed wavy trajectories. We use the geometry shown in Fig. 4b and described in the text. Typical Reynolds numbers for swimming *Gonium* are ~ 10^−3^, in Stokes regime. The velocity of the surrounding fluid at a position x can therefore be expressed as boundary integrals,

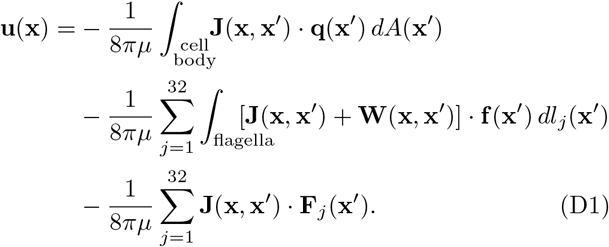

Here, **J** is the Stokeslet kernel, **W** the kernel for slender body theory, **q** the traction force on the cell body, **f** the traction force on flagella, and **F** the point force generated by the flagella. The first term of the r.h.s of Eq. (D1) represents the drag of the cell body, which is solved by a boundary element method with a mesh of 320 triangles. The second term accounts for the drag of the 32 flagella, which is solved by slender body theory with 10 slender elements per flagellum. The third term represents thrust and spin forces generated by flagella. A no-slip velocity boundary condition is applied on the cell body and along the flagella; force- and torque-free conditions are applied to the whole body. Detailed functional forms and the velocity along the flagella can be found in Itoh *et al*.[33]; here we solve Eq. (D1) in a similar manner.

The amplitude *A* of the oscillations of the computed wavy trajectories, using the force variation around *Gonium* given by Eq. (5), linearly increases with the strength *ξ* of the imbalance (Fig. 12), while the wavelength λ is hardly impacted. The deduced pitch angle *χ* therefore also linearly increases with *ξ*, in good agreement with Eq. (6), plotted by the dashed line. This enables us to deduce the experimental imbalance amplitude, as *χ* ≃ 30° corresponds to *ξ* ≃ 0.4. Finally, the decrease in swimming efficiency *v_m_*/*v* follows the geometrical prediction *v_m_*/*v* = cos *χ*, as plotted in Fig. 3c.

**FIG. 12.**
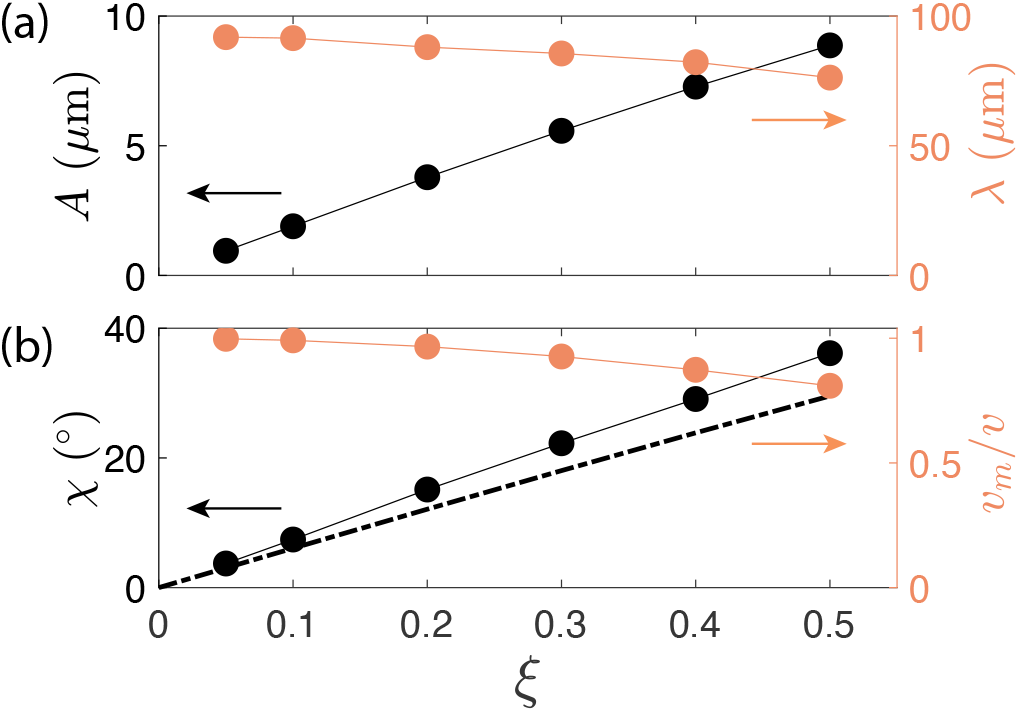
Properties of wavy helical trajectories. (a) Amplitude *A* of the oscillations and wavelength λ of the helix as a function of the defect parameter *ξ*. (b) Pitch angle *χ* and swimming efficiency *v_m_*/*v* as a function of the defect parameter *ξ*. The dashed line shows the theoretical prediction for *χ* as a function of *ξ*, obtained from Eq. (6).

## Appendix E: Maximum of the adaptive response

Defining *s* = *t*/*τ_r_* in the step response (9) for *t* > 0 we have

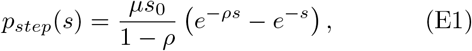

where again *ρ* = *τ_r_*/*τ_a_*. The maximum response obtained by setting *dp_step_*/*ds* = 0 occurs at *s*\* = −(1 − *ρ*)^−1^ ln *ρ*, with the magnitude 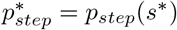 given by the amusing function

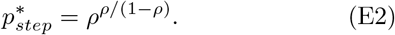

As shown in Fig. 13, this has a maximum at *ρ* = 0, confirming the statement in Sec. IIIC that the maximum amplitude response is found in the limit *τ_r_*/*τ_a_* → 0.

**FIG. 13.**
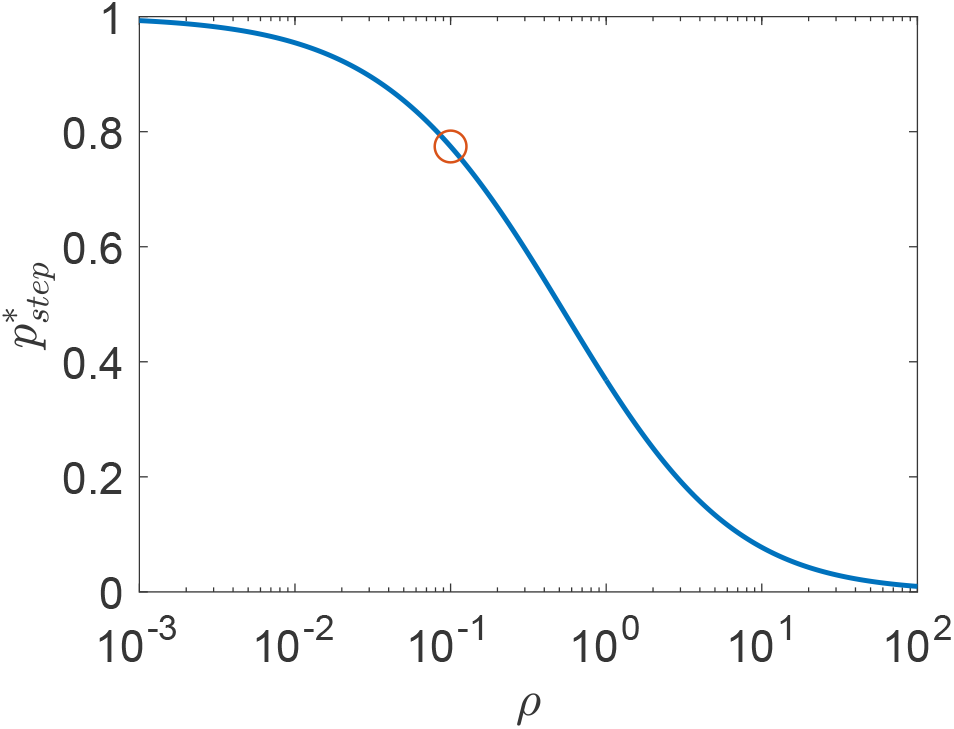
Maximum amplitude (E2) of the adaptive phototactic response to a step-up in light intensity, as a function of the ratio *ρ* = *τ_r_*/*τ_a_*. Red circle indicates experimental average.

## Appendix F: Phototactic gain function

We estimate here the photoresponse *p_α_* at an arbitrary angle *α* around *Gonium*. This allows computation of the phototactic torque (3) from the flagella force (7). The balance with the viscous torque eventually leads to the phototactic reorientation trajectory. This implies the definition of a gain function 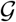. From (10), the response to light stimulation *s*(*t*) is

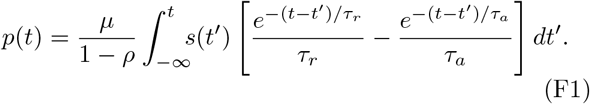

In this model, *p*(*t*) has a slight negative overshoot when *s*(*t*) decreases. However, for *τ_r_* ≪ *τ_a_*, one has *p*(*t*) > 0 for almost all time, and the integration of *p* among the flagella on the illuminated side remains positive.

To evaluate the integral, we use the following geometry. We consider that only flagella on the illuminated side of the *Gonium*, defined by *α* = −*π*/2 ⋯ *π*/2, contribute to the reorientation torque. A flagellum at angle α has traveled from −*π*/2 to *α* at constant spin *ω*_3_. After its transit to the dark side, each flagellum looses the memory of its previous transit on the illuminated side, so that *p* = 0 when it reaches *α* = −*π*/2. With *s*(*α*) = *s*_0_ cos *ϕ*cos *α* and using *α* = *ω*_3_*t*, Eq. (F1) becomes

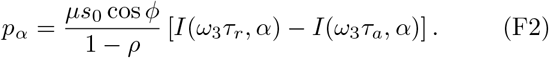

with

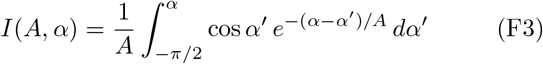

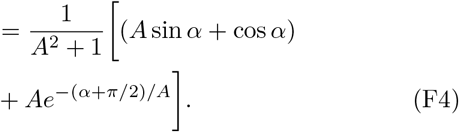

We finally compute the torque (3) from the flagella force (7). Because of the delay induced by the adaptive response, the torque is no longer along **ê**_*z*_, implying a change in *θ*. Since we assumed *θ* = *π*/2 earlier, we consistently neglect this effect, and consider only the *z* component of the torque,

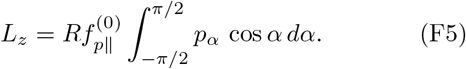

Inserting (F2), we rewrite this integral as

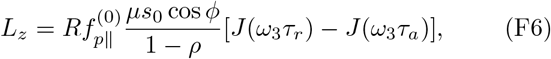

where

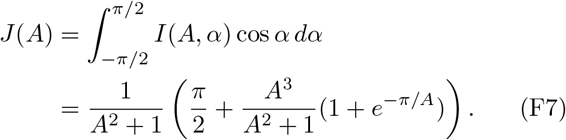

We have *J*(0) = *π*/2 and *J*(*A*) ≃ 2/*A* for *A* ≫ 1. Balancing this torque with the *z* component of the friction torque 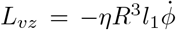 yields a differential equation in the form (14), from which we identify the gain function

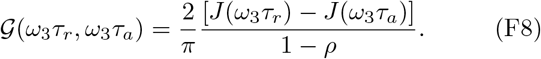

